# The DeepFaune initiative: a collaborative effort towards the automatic identification of French fauna in camera-trap images

**DOI:** 10.1101/2022.03.15.484324

**Authors:** Noa Rigoudy, Gaspard Dussert, Abdelbaki Benyoub, Aurélien Besnard, Carole Birck, Jérome Boyer, Yoann Bollet, Yoann Bunz, Gérard Caussimont, Elias Chetouane, Jules Chiffard Carriburu, Pierre Cornette, Anne Delestrade, Nina De Backer, Lucie Dispan, Maden Le Barh, Jeanne Duhayer, Jean-François Elder, Jean-Baptiste Fanjul, Jocelyn Fonderflick, Nicolas Froustey, Mathieu Garel, William Gaudry, Agathe Gérard, Olivier Gimenez, Arzhela Hemery, Audrey Hemon, Jean-Michel Jullien, Daniel Knitter, Isabelle Malafosse, Mircea Marginean, Louise Ménard, Alice Ouvrier, Gwennaelle Pariset, Vincent Prunet, Julien Rabault, Malory Randon, Yann Raulet, Antoine Régnier, Romain Ribière, Jean-Claude Ricci, Sandrine Ruette, Yann Schneylin, Jérôme Sentilles, Nathalie Siefert, Bethany Smith, Guillaume Terpereau, Pierrick Touchet, Wilfried Thuiller, Antonio Uzal, Valentin Vautrain, Ruppert Vimal, Julian Weber, Bruno Spataro, Vincent Miele, Simon Chamaillé-Jammes

**Affiliations:** CEFE, Université de Montpellier, CNRS, EPHE, IRD, Montpellier, France; Université Lyon 1, CNRS, UMR 5558, Laboratoire de Biométrie et Biologie Evolutive, Villeurbanne, France; Parc National des Ecrins, Domaine de Charance, Gap, France; Asters, Conservatoire d’espaces naturels de Haute-Savoie, 84 route du Viéran, PAE de Pré Mairy, Pringy, France; Office Français de la Biodiversité, Direction de la Recherche et Appui Scientifique Micropolis, Gap, France; Fédération Départementale des Chasseurs de l’Ain, 19 Rue du 4 Septembre, Bourg-en-Bresse, France; Fond d’Intervention Eco-Pastoral (FIEP) Groupe Ours Pyrénées, 1 rue Boyrie, Pau, France; Université Grenoble Alpes, CNRS, Université Savoie Mont Blanc, LECA, Laboratoire d’Ecologie Alpine, Grenoble, France; Observatoire spatio-temporel de la biodiversité et du fonctionnement des socio-écosystèmes de montagne (ORCHAMP), France; Office Français de la Biodiversité, Direction de la Recherche et Appui Scientifique, Service “Santé de la faune et fonctionnement des écosystèmes agricoles”, France; Institut du Développement et des Ressources en Informatique Scientifique (IDRIS), IDRIS-CNRS Campus universitaire d’Orsay, rue John Von Neumann, Orsay, France; Centre de Recherches sur les Ecosystèmes d’Altitude (CREA Mont-Blanc), Observatoire du Mont-Blanc, Chamonix, France; Parc naturel régional du Perche, Courboyer, Perche-en-Nocé, France; CERFE - URCA, 5 rue de la Héronnière, Boult-aux-Bois, France; Parc naturel régional des Marais du Cotentin et du Bessin, 3 village Ponts d’Ouve, Saint-Côme-du-Mont, Carentan-les-Marais, France; Fédération Départementale des Chasseurs du Jura, Route de la Fontaine Salée, Arlay, France; Parc national des Cévennes, 6 bis, Place du palais, 48400 Florac-Trois-Rivières, France; Institut Méditerranéen du Patrimoine Cynégétique et Faunistique, Domaine expérimental agri-environnement, Villa “Les Bouillens”, Vergèze, France; Office Français de la Biodiversité, Direction de la Recherche et de l’Appui Scientifique, Montfort 01330 Birieux & 5 allée Bethleem Gières, France; Parc naturel régional des Ballons des Vosges, gestionnaire de la RNN de la Tourbière de Mâchais, 1 place des verriers, Wildenstein, France; Etablissement public du Mont-Saint-Michel, 16 route de la Caserne, Beauvoir, France; Chemin de Pré Rond-Les Mariages, Le Châtelard, France; OEKO-LOG field research Joachimsthaler Str. 9, 16247 Parlow, Germany; Fauna & Flora International, The David Attenborough Building, Pembroke Street, Cambridge, CB2 3QZ, United Kingdom; Parc Naturel Regional des Baronnies Provençales, 575 route de Nyons, 26510 Sahune, France; GEODE UMR 5602, CNRS, Université Jean-Jaurès, 5 Allée Antonio-Machado, 31058 Toulouse, France; Parc national du Mercantour, 23 rue d’Italie CS 51316 06006 Nice, France; Institut de Recherche en Informatique de Toulouse, DPD - Université Toulouse III - Paul Sabatier, 118 Route de Narbonne, Toulouse, France; Fédération Départementale des Chasseurs de la Drôme, 3132 Route des Sétérées, Crest, France; 100 rue Barbara 34730 Prades le Lez, France; Parc naturel régional de l’Aubrac, Place d’Aubrac, Aubrac, France; 2 la Place du Château, 34190 Brissac, France; Office Français de la Biodiversité, Direction de la Recherche et Appui Scientifique, Service “Conservation et Gestion des Espèces à Enjeux”, France; School of Animal, Rural and Environmental Sciences, Nottingham Trent University, NG25 0QF, United Kingdom; Office Français de la Biodiversité, Service Départemental, 12000 Rodez, France

**Author notes:** Contributed equally (Noa Rigoudy, Gaspard Dussert).

**Keywords:** camera traps, classification, deep-learning, Europe, France, software

## Abstract

Camera traps have revolutionized how ecologists monitor wildlife, but their full potential is realized only when the hundreds of thousands of collected images can be readily classified with minimal human intervention. Deep-learning classification models have allowed extraordinary progress towards this end, but trained models remain rare and are only now emerging for European fauna. We report on the first milestone of the DeepFaune initiative (https://www.deepfaune.cnrs.fr), a large-scale collaboration between more than 50 partners involved in wildlife research, conservation and management in France. We developed a classification model trained to recognize 26 species or higher-level taxa. The classification model achieved 0.97 validation accuracy and often >0.95 precision and recall for many classes. These performances were generally higher than 0.90 when tested on independent out-of-sample datasets for which we used image redundancy contained in sequence of images. We implemented our model in a software to classify images stored locally on a personal computer, so as to provide a free, user-friendly and high-performance tool for wildlife practitioners to automatically classify camera-trap images.

## 1. Introduction

Camera traps have revolutionized the way ecologists monitor biodiversity and population abundances (O’Connell et al. 2011; Howe et al. 2017). Camera traps enable to scale-up monitoring efforts dramatically, both in space and time, as they are relatively cheap, easy to deploy and autonomous tools (Steenweg et al. 2017). The continuous monitoring capacity they provide also facilitates the detection of rare species.

Therefore, and unsurprisingly, it has become common practice to deploy tens to hundreds of camera traps in monitoring programs.

The full potential of camera traps is however only realized when the hundreds of thousands of images, many being empty from spurious detections, can be rapidly classified with minimal human intervention (Chen et al. 2014; Schneider et al. 2018; Wearn et al. 2019; Tuia et al. 2022) Since the beginning, machine learning approaches, and in particular deep learning models, have held the promise to solve this issue. Recent works have confirmed their power: for instance, deep-learning models developed by different research teams (Willi et al. 2019; Tabak et al. 2019; Whytock et al. 2021) have obtained > 0.90 recall and precision for a number of mammal species in North American ecosystems, African savannas and tropical forests. The recurring leading approach, as assessed in recent iWildcam competitions (Beery et al. 2021), consists in two steps: (step 1) detecting animals, humans and vehicles and filtering out empty images using a robust detection model such as MegaDetector (https://github.com/microsoft/CameraTraps/; Beery et al. 2019) and (step 2) using a CNN classification model to identify the species in the images when an animal was detected. While MegaDetector is efficient in detecting animals in various environment and geographical regions, there is still a need for tailored CNN classification models dedicated to the species and regions of interest.

The number of image classification models available to ecologists currently remains low and their taxonomic coverage is still limited. So far, most of the image classification models developed on camera-trap data have been trained, even when based on millions of pictures, using images collected by a few partners in one or two sites. For example, Norouzzadeh and colleagues (2018) trained their model on 3.2 million images taken from the Serengeti Snapshot project (Swanson et al. 2015) to identify 48 African mammals, yet all the data was collected from one large study area in the Serengeti National Park as part of the same monitoring project. Although this can lead to highly accurate models when tested on data collected in the same sites or applied to similar systems, such models may fall short from generalizing to other fauna or contexts (Beery et al. 2018). For the European fauna, we believe there is potential in aggregating multiple small datasets from a large number of partners to build sufficiently large training, validation and test datasets to develop efficient classification models. The pros and cons of a cross-partner image aggregation strategy, and its ability to lead to successful models, have yet to be investigated. Recent initiatives such as Wildlife Insights (www.wildlifeinsights.org) or Agouti (www.agouti.eu), online platforms to manage and classify camera-trap images, have embraced this strategy. Naturally, each initiative will have its own terms of participation, which may not be acceptable for certain institutional partners. For example, online platforms require the users to upload their images, and sometimes have non-optional data-sharing policies. This approach may not be suitable for certain institutional partners which have legally-bounding contracts on data collection or privacy concerns. Some other initiatives offer users solutions to run models on their own servers or computers (Trapper: https://trapper-project.readthedocs.io; Project Zamba: https://zamba.drivendata.org; our work).

Developing a classification model is however not only a question of collecting a large amount of data. Many issues appear when going beyond proof-of-concept and aiming at propose a model that should be of high-enough quality in practice (e.g. data leakage, lack of transferability, shortcut learning, sensitivity to background, out-of-domain predictions, calibration, unbalanced data and biaises for rare species; Desprez et al. 2023). Therefore, we advocate that releases of species-classification models for the ecological community must be accompanied by a reliable performance analysis to help ecologists understand how well the proposed classification model is expected to perform in practical situations (e.g. on new camera traps images).

We report here on the DeepFaune initiative (https://www.deepfaune.cnrs.fr). This initiative aims at (1) aggregating camera-trap images from many French institutions or research groups to create a collective and large-scale dataset, (2) developing a deep-learning-based species classification model, using an clearly explained methodology to circumvent the issues mentioned previously, (3) performing a rigorous performance analysis on out-of-sample datasets, (4) releasing the model to the community as a software component that can be integrated in other tools and (5) providing a free, open-source, user-friendly and efficient software for practitioners to run the identification pipeline on self-hosted images with standard personal computers.

## 2. Methods

### 2.1 Partners and data collection

At the time of writing, the initiative brings together over 50 partners (see the complete list at https://www.deepfaune.cnrs.fr) representing institutions managing protected areas, hunting federations or academic research groups across France. A few partners also joined from other European countries. Partners provided camera-trap images or videos, originating from various monitoring projects (e.g. general monitoring of Alpine fauna, monitoring of the burrows of badgers) or opportunistic surveillance (e.g. to assess the presence of wolves). Among these partners, five partners joined the project after initial data collection, providing annotated pictures to use as out-of-sample data for a stringent test of the classification model accuracy (see below).

### 2.2 Original dataset

We gathered and sorted more than 2 millions annotated pictures and 20 thousands annotated videos, after removal of corrupted or other problematic files. These pictures and videos included either one of the animal species originally identified by the partners, a human, a vehicle, or were empty (less than 1%), i.e. had no animal, human or vehicle visible. In some instances, annotations had been made at a higher taxonomic level than the species level (e.g. bird, rodent). The amount of data provided by each partner varied a lot, ranging from 20 videos to hundreds of thousands of images. Many partners shared data from camera traps set to take a series of images for a single detection event. We thereafter defined these consecutive images as ‘sequences’, considering that two images taken within 10s, at the same site, belonged to the same sequence. Finally, as we wanted to train our model to classify camera-trap images, we converted videos into images by retaining one frame per second out of the first 4 seconds of each video.

### 2.3 Targeting 26 animal species or higher-order taxonomical groups

We restricted our dataset and model to 26 animal species or higher-order taxonomical groups that were common enough and well sampled in our original dataset: European badger (*Meles meles)*, bear (Ursus arctos), bird, chamois (alpine, *Rupicapra rupicapra*, and Pyrenean, *Rupicapra pyrenaica*), cat (domestic *Felis catus* and wild), cow (*Bos Taurus)*, dog (*Canis familiaris)*, equidae, red fox (*Vulpes vulpes*), common genet (*Genetta genetta*), goat (*Capra* sp.), hedgehog (*Erinaceus europaeus*), ibex (*Capra ibex*, alpine only), lagomorph, lynx (*Felix lynx)*, marmot (*Marmota marmota)*, micromammal, mustelidae, mouflon (*Ovis gmelini)*, nutria (*Myocastor coypus*), red deer (*Cervus elaphus)*, roe deer (*Capreolus capreolus)*, sheep (*Ovis aries*), squirrel (*Sciurus vulgaris)*, wild boar (*Sus scrofa)* and wolf (*Canis lupus lupus*).

### 2.4 Training and validation datasets

#### Avoiding shortcut learning

We prepared the training and validation datasets using MegaDetectorV5 (model v5a; (Beery et al. 2019) on our original dataset to filter out empty images and images containing humans and/or vehicles. MegaDetector v5a is based on the image detection model YOLOv5 (Redmon et al. 2016) and allowed us to produce one cropped image per animal that was detected (there were potentially multiple individuals in a given image). We retained cropped images that had a confidence score higher than 0.8 to decrease the risk of false detection. Since the images in the original dataset were annotated, it was possible to annotate any cropped image in the training and validation datasets using the annotation of its parent image. For instance, if MegaDetectorV5 detected two animals in an image originally annotated as ‘wild boar’, we considered that the two resulting cropped images contained a wild boar to train the model. After this step, we remained with a dataset of 787 575 cropped animal images.

In addition to often having, for most species, images taken in dozens of different settings thanks to the numerous contributors, the ‘cropping’ step described above also enabled us to avoid shortcut learning (Geirhos et al. 2020) as we trained and validated our CNN classification model on annotated images whose background had been mostly removed.

### Avoiding data leakage

CNN models require independent training and validation datasets to learn successfully (Kapoor and Narayanan 2022), and as commonly done, we split the whole image dataset into a training dataset containing 90 % of the images, and a validation dataset containing the remaining 10%. One issue that could arise during this split is that pictures collected within the same sequence are used in both the training and validation datasets, which would threaten their independence (Desprez et al. 2023). As our original dataset contained pictures and videos collected with many different camera-trap models, there was no consistency in how the date and time stamps were stored. We thus chose to identify sequences through image filenames instead. We used a heuristic based on text mining in the R package *stringdist* (van der Loo 2014) to compare filenames to each other. We considered that two files were not independent if their filenames were too similar, i.e. if their similarity was above 0.9. This conservative threshold insured there was no image from the same sequence or video that was present in both the training and the validation datasets.

### Dealing with class imbalance

Class imbalance, that is the fact that the number of images per class differs, is also known to affect both training and validation (Johnson and Khoshgoftaar 2019). Here, we propose a novel approach that combines downsampling and upsampling during the training and validation phases. At each epoch of the training phase, we downsampled the class which was overrepresented down to a multiple of the rarest class. We chose to have a ratio of 5 between the number of images of the most common class and the number of images of the rarest class. This was estimated for every epoch, such that every image is seen by the model if we fit the model for enough epochs. As a consequence, images of the rarest classes were used in many epochs and can thus be considered upsampled whereas those of the most common classes are downsampled. This way, the optimization problem was different at every epoch, but more balanced. We considered that the stochasticity induced by the sampling process at each epoch had a positive impact on the model fit procedure and we observed a continuous reduction in model loss through epochs. We used the same idea to handle imbalance during validation. This enabled us to compute what we called a balanced validation accuracy.

### 2.5 Training and validation phases

We used transfer learning starting from a ConvNext-Base model (Zhuang et al. 2022) that was pre-trained on Imagenet 22K, with a resolution of 224 × 224, using PyTorch. We performed image augmentation using the *imgaug* Python library (https://github.com/aleju/imgaug) during the training phase. Each image in each batch was modified with a random set of transformations such as horizontal flip, affine transformations, gray scale transformation, blurring, brightness or contrast changes. We used the softmax output as prediction score. We used the SGD optimizer with a learning rate of 4e-4. We then estimated the model for a maximum of 100 epochs, monitored the overall balanced validation accuracy and stopped the estimation when it did not increase with a patience of 10 epochs.

For each processed image, the model computes a prediction score for each class (these scores sum to 1.). The prediction score expresses the certainty on the attribution of the image to a given class. In the following, we considered as a prediction the class maximizing these score. To estimate the quality of the model, we computed global accuracy (i.e. the probability that a prediction is correct), precision (i.e. the probability that an image classified as showing species i actually has species i present) and recall (the true positive rate, i.e. the probability that an image having species i present is actually classified as showing species i).

### 2.6 Estimating classification performance on out-of-sample data using sequences

#### Building an out-of-sample dataset

We investigated the ability of the pipeline (i.e. the detection model followed by the classification model) to perform an accurate identification of species in an out-of-sample dataset, i.e. on images taken in contexts that have never been seen during the training stage (Schneider et al. 2020). Our out-of-sample dataset originated from five partners plus a public dataset from the Peneda-Gerês National Park in Portugal (Zuleger et al. 2023). We explored the classification accuracy of the CNN-based classification step after using MegaDetector V5 (model v5a) to retrieve cropped images around animals using a threshold of 0.8. The final out-of-sample contained 213 059 cropped animal images of 23 of the 26 species or higher taxonomic groups on which the Deepfaune pipeline focuses (lynx, hedgehog and nutria were lacking).

### Using sequence information to enhance prediction

We predicted the species present at the level of the sequence and not at the level of the individual image. We remained with a out-of-sample dataset of 36 033 sequences. We chose to predict on the sequence level for two reasons: camera traps are now often set to take images in bursts, and animals commonly stay several seconds near the camera, triggering it multiple times in a row. To do so, we extracted the date and time of each image of the out-of-sample dataset using the EXIF information. We obtained the prediction score at the sequence level by first computing the prediction score for each image of the sequence that was not predicted as empty. We then summed these scores and considered that the class with the highest total prediction score was the predicted class for the sequence. Images that were not in a sequence were classified using their own prediction score. Sequences with a score lower than 0.5 were considered as ‘undefined’.

### 2.7 The DeepFaune software

We found that MegaDetectorV5 created a bottleneck in terms of speed when used on a CPU (about 2 to 3 seconds per image in our experiments on various personal computers) due to its high image resolution (1280) and large model size based on YOLOv5x (https://github.com/ultralytics/ultralytics). We therefore developed a much faster alternative using YOLOv8s (https://github.com/ultralytics/ultralytics), which is a high-performance medium-sized detection model with a resolution of 640. We trained the YOLOv8s-based detector using the cropping information given by MegaDetectorV5 in images previously used in this study (to limit the computing needs, we took a subset of our full dataset). We obtained a detection performance that is generally similar to the one of MegaDetectorV5, but at a much higher speed (see Online Resource 1 Table S.1). We then implemented this alternative detector (which needs only about 0.3 second per image) and our CNN classifier inside a free, user-friendly and independent software. This enables users to input their images, stored locally, directly in the software, and classify these images either as empty, human, vehicle or one of the 26 animal classes on which the CNN classifier was trained. Users can decide of a threshold on the prediction score below which the images will not be classified and will be set aside in a ‘undefined’ class for visual inspection. The software only requires the installation of Python v3 and its PyTorch module. To overcome the common difficulties associated with the installation of Python libraries on Windows operating systems, we created a standalone and ready-to-use .exe program for these systems. The software is available on the project’s website: https://www.deepfaune.cnrs.fr.

## 3 Results

### 3.1 Performance of the classification model on the validation dataset

We obtained an overall balanced validation accuracy of 97.3%. Recall and precision were >0.9 for most classes (Table 1), and notably above 0.95 for many classes for which the number of training images was high (e.g. cow or wild boar). Interestingly, we obtained high performance scores for lynx regardless of the small training sample size (806 images) for this species. This may be due to its characteristic coat color and spotted pattern which facilitates recognition and classification.

**Table 1.** Performance metrics of the classification model computed on the validation dataset. N Training and N Validation refer to the number of images of a given class within the training and validation datasets, respectively.

| Class | N Training | N Validation | Precision | Recall |
| --- | --- | --- | --- | --- |
| badger | 9263 | 1227 | 0.96 | 0.98 |
| ibex | 6168 | 756 | 0.83 | 0.97 |
| red deer | 130753 | 11468 | 0.96 | 0.96 |
| chamois | 113527 | 14465 | 0.99 | 0.96 |
| cat | 5013 | 926 | 0.90 | 0.94 |
| goat | 3459 | 235 | 0.83 | 0.93 |
| roe deer | 81923 | 8167 | 0.95 | 0.95 |
| dog | 9413 | 1202 | 0.94 | 0.93 |
| squirrel | 4771 | 576 | 0.94 | 0.98 |
| equid | 1211 | 80 | 0.91 | 0.99 |
| genet | 340 | 52 | 0.89 | 0.92 |
| hedgehog | 231 | 2 | 0.50 | 1.00 |
| lagomorph | 4655 | 764 | 0.94 | 0.96 |
| wolf | 7096 | 959 | 0.87 | 0.96 |
| lynx | 806 | 89 | 0.95 | 1.00 |
| marmot | 1934 | 228 | 0.95 | 0.98 |
| micromammal | 3265 | 444 | 0.99 | 1.00 |
| mouflon | 967 | 72 | 0.60 | 0.96 |
| sheep | 113691 | 15566 | 1.00 | 0.99 |
| mustelid | 4515 | 619 | 0.87 | 0.95 |
| bird | 26575 | 4069 | 0.99 | 0.99 |
| bear | 6672 | 1362 | 0.91 | 0.98 |
| nutria | 382 | 63 | 0.94 | 0.97 |
| fox | 47064 | 4558 | 0.98 | 0.95 |
| wild boar | 70616 | 13300 | 0.98 | 0.99 |
| cow | 44773 | 7241 | 0.99 | 1.00 |

We did not observed many identification mismatches between the different ungulate species (Online Resource 1, Fig. S.1). For instance, performance for red deer and roe deer was excellent, despite some objective visual similarities. Precision for mouflon was not great (0.72), but this was because of the unbalance in the validation set: since there was at least twice as many chamois or sheep as mouflon images, having only a few of the chamois and sheep misclassified as mouflon was enough to lead to a low precision for mouflon. The same reason explained the low precision for goat (0.83). Surprisingly, we obtained a moderate performance for ibex: indeed, mismatches appeared on images where ibex’s distinctive horns were not visible on the images. Finally, we observed a lower-than-desired precision for wolf. This was due to some confusion between wolf and fox and dog, mostly on night images in which species identification was intrinsically challenging. Again, this precision is also driven by the aforementioned unbalance in the validation set.

### 3.2 Performance of the classification model on the out-of-sample dataset

The classification model achieved an overall accuracy of 93.6% on individual images and 96.7% on the out-of-sample dataset when predicting at the level of image sequences. Class-specific results were also generally significantly better when identification was done at the sequence level (Table 2). For instance, the recall for roe deer increased from 0.92 to 0.96 using sequences. This suggested that fine-grained species identification was facilitated by having different views of the same moving animal (see an example in Fig. 1). The performance using sequences are overall close to those obtained on the validation dataset (without sequences).

**Table 2.**
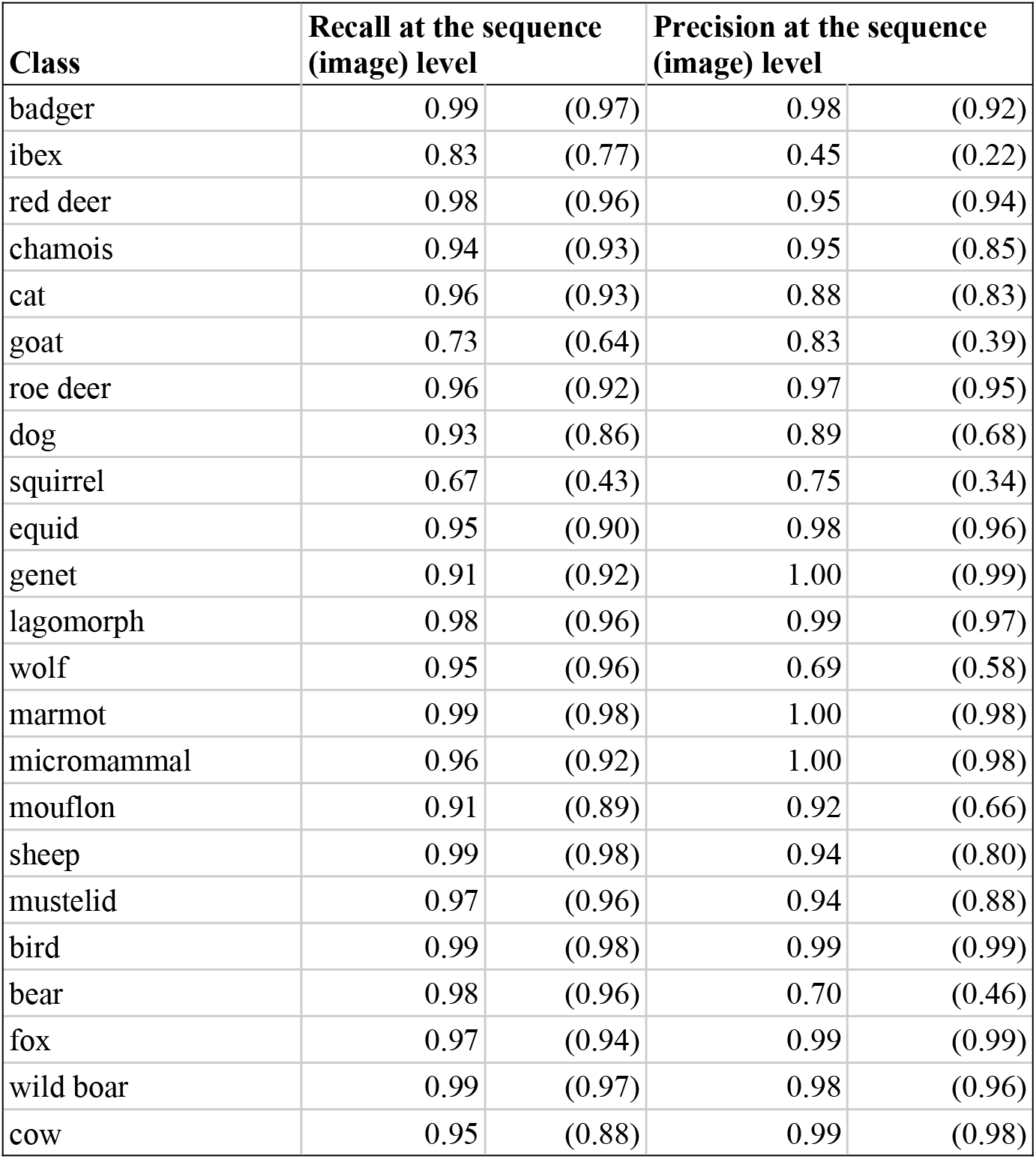
Performance metrics of the classification model computed on out-of-sample data, when predicting at the level of individual images or image sequences. Model performance at the image level is expected to be lower than at the sequence level as people who annotated the images had the complete 3-images sequences available and could use previous or subsequent images to clarify uncertainties. Images and sequences that had a prediction score lower than 0.5 where not considered and classified as ‘undefined’ (4.07% and 5.59% of the images and sequences, respectively).

**Figure 1.**
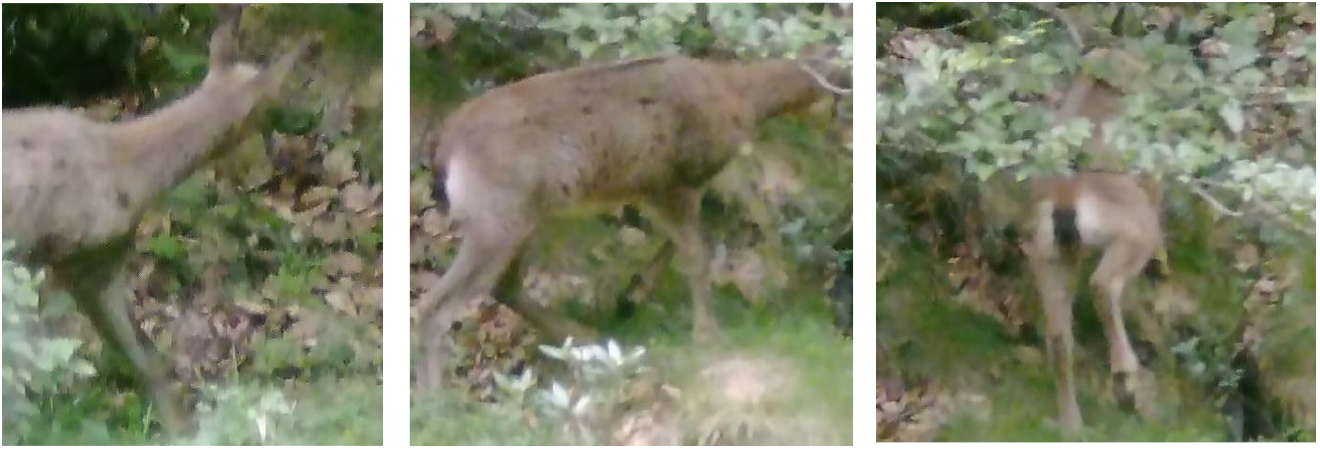
Using sequences decreases the false positive rate in classification predictions. A sequence of three images of a chamois, taken over a 3-second interval (and cropped by MegaDetectorV5). The two first images are predicted as chamois (with scores 0.91 and 0.96). The third image is misclassified as a mouflon with a lower score (0.81), but the score at the sequence-level is highest for chamois. The three images are therefore predicted to show a chamois by the DeepFaune pipeline.

High performance was achieved for large ungulates, with precision and recall being about 0.95 for red deer, roe deer, chamois, and wild boar (Table 2), with the exception of ibex and mouflon for which the model made some confusion with other species (e.g. goat). There remained some mismatches between classes (Online Resource 1 Fig. S.2), for instance 155 sequences of roe deer that were classified as red deer.

Performance remained excellent for smaller animals, including marmots, lagomorphs, micromammals and birds. Contrary to what is commonly expected, we observed almost no confusion between wolf and dog predictions, which suggested that our model could distinguish well between the two. As in the validation dataset, inspection of the confusions revealed that they mostly occurred mostly with nighttime images or when the animal was partially visible (see examples in Fig. 2).

**Figure 2.**
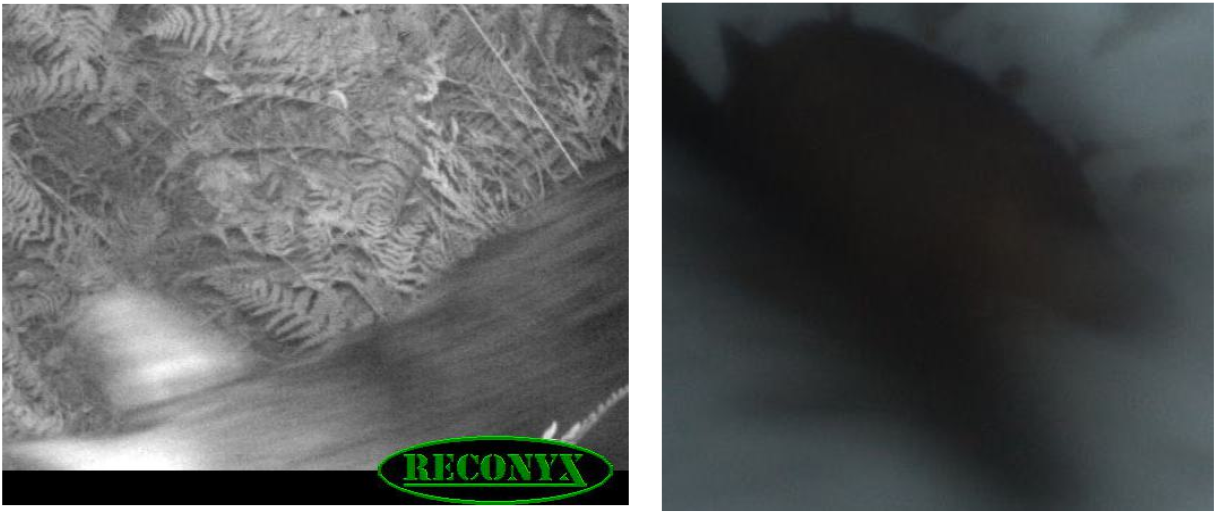
Examples of images in which a wolf is not correctly identified, possibly due to poor image quality. A wolf was classified as a roe deer, with a low prediction score of 0.51 in a sequence with only one non-empty image (left), and as a fox with a moderate score of 0.82 in a sequence of 1 image (right). Both images were cropped by MegaDetectorV5.

### 3.3 Thresholding prediction scores to balance error rate vs. increased manual labeling effort

Most of the prediction errors are associated with a moderate to low or very low score (see Fig. 2 for example). To improve the likelihood that species are correctly predicted by the classification model, one can therefore consider as correct only predictions that have a score above a certain threshold, and change the predictions that have a lower score to an ‘undefined’ class. Images with predictions being ‘undefined’ would have to be inspected manually to be correctly labelled. As the score for threshold predictions is increased, the number of errors is reduced, but the number of images to be annotated manually increases. Fig. 3 shows how these numbers vary with the score threshold for the classification model presented in this work. In the DeepFaune software, we use a score threshold of 0.8 by default, leading to an error rate of less than 1% on the out-of-sample dataset, at the cost of having to manually annotate approximately 30 % of the images. This score threshold can be modified by the user.

**Figure 3.**
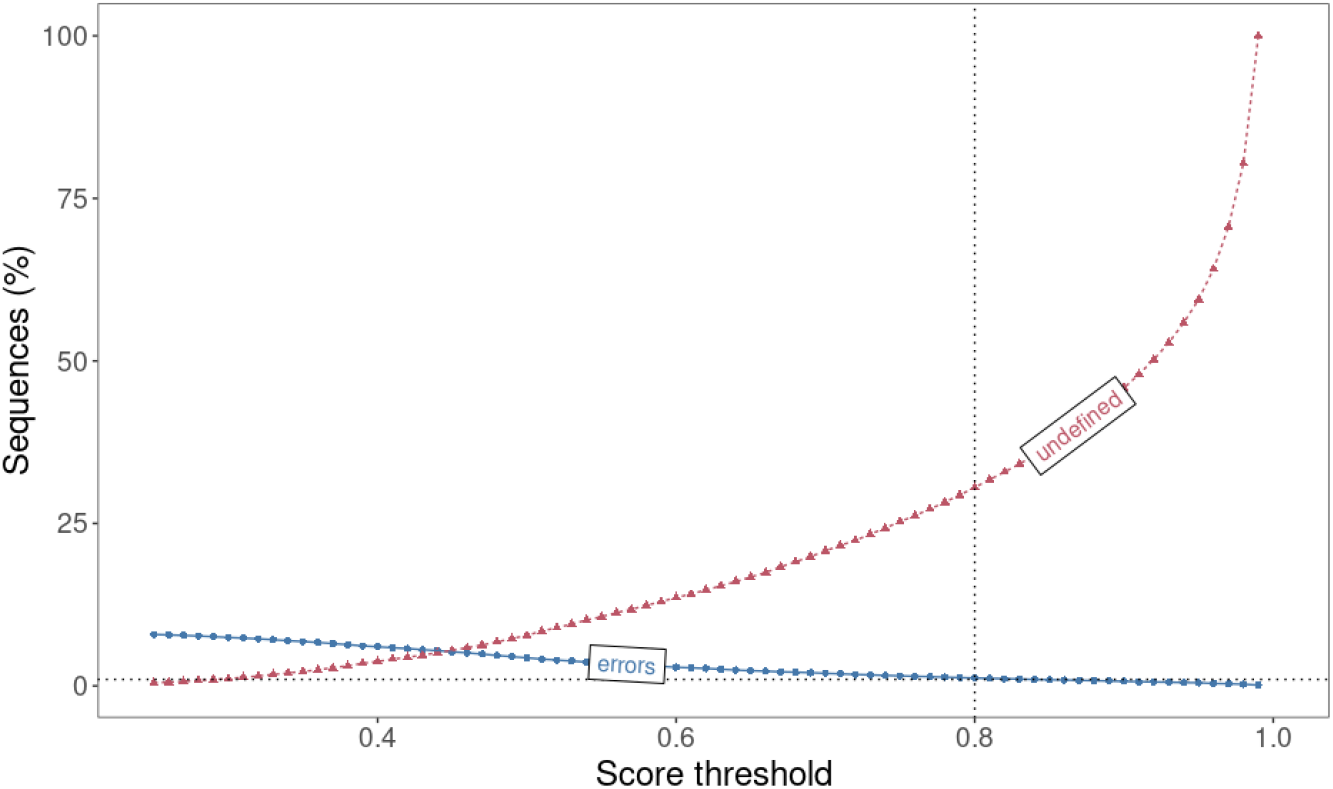
Effect of increasing the prediction score threshold value on the percentage of image sequences that are wrongly classified (‘errors’; in blue squares) or classified as ‘undefined’ (red triangles). The analysis is conducted on the out-of-sample dataset. The vertical dotted line is at 0.8, the default score threshold value used in the DeepFaune software. The horizontal dotted line indicates 1% of the sequences.

## 4 Discussion

The DeepFaune initiative represents a successful multi-partner collaboration to build a large-scale camera-trap image dataset and develop a machine learning model, fully tested and evaluated, readily available to automatically classify camera-trap images collected in Europe. Our current model allows for predicting the presence of 26 different species (of higher order taxa) in camera-trap images. Although we did use the standard approach of transfer learning, we implemented a number of tricks to deal with class imbalance and the independence of training and validation datasets that could be useful to others. Ultimately, our species classification model performed extremely well on the validation dataset and provided robust results on out-of-sample data. Additionally, we built a graphical user interface (GUI) that is freely available online (https://www.deepfaune.cnrs.fr) so that practitioners can easily run the model locally on a personal computer and automatically sort their images or videos based on the model predictions. The model is also available as a software component to be integrated in other tools, in particular camera-trap management online platforms, such as Agouti (www.agouti.eu) for instance, in which the Deepfaune classification model can already be used.

### 4.1 Model performance and relevance for ecological studies

There are currently few species-recognition models available for the European fauna and our results could therefore be used to benchmark future studies. The quality of our model appears very good, even by deep-learning classification standards, and compares favorably with results from similar exercises conducted on fauna from other continents. For instance, overall accuracy of 0.97, 0.78 and 0.94 on validation datasets were respectively reported in studies on North American fauna (Tabak et al. 2019), Central African fauna (Whytock et al. 2021) and East African fauna (Norouzzadeh et al. 2018). Our model also perform better than those reported by Carl et al. (2020) and Simões et al. (2023) that are the rare existing models focused on European fauna and for which performance metrics are available.

We managed to obtain good results with a moderately sized dataset (by deep-learning standards), by overcoming a number of obstacles. Firstly, the large imbalance (e.g. two orders of magnitude more images of chamois than ibex) would have biased the classification towards the most common species. We therefore implemented an original approach combining downsampling and upsampling techniques that successfully prevented the emergence of a relationship between identification performance and the number of images per class. Secondly, deep learning models are known to be able to learn ‘shortcuts’ (Geirhos et al. 2020) and can misclassify images due to spurious similarities (e.g. similar image backgrounds). In our context, this would correspond to learning contextual elements (e.g. snow, peculiar vegetation) that would be associated with the species, rather than learning to recognize the species itself. To avoid this potential issue, we chose to classify cropped images, and not whole images (as opposed to Whytock et al. 2021), with an additional procedure of image augmentation.

Overall, class-specific recall and precision are high enough for the model to be useful for many specific studies, some of which we highlight now: (1) automatized monitoring of large ungulates, a guild of important management interest in Europe. Ungulates are indeed generally very well classified by the model, with recall and precision values above 0.9 for most species (as explained in the Results section, performance for ibex and mouflon are low, but will likely improve as new pictures are continuously added in the database). These results suggest that our model could be a useful tool to facilitate studies investigating the effects of management practices on locally abundant ungulates (e.g. can wild boar population dynamics be controlled by hunting?) or the dynamics of prey under predation (e.g. do roe deer populations decrease as wolves return?); (2) monitoring large mammalian communities. Although model accuracy levels were generally lower for classes other than ungulates, the model’s performance and taxonomic coverage suggest that it could be useful to automatically sort out hundreds of thousands of pictures for a range of taxonomic groups. This could prove especially useful in studies looking at the impact of anthropization on large mammal communities for example. A human intervention would still be required to manually verify the pictures for which prediction scores are low or to identify the species present in images where the model classifies at a higher taxonomic level (e.g. mustelidae), but this effort should be dramatically reduced when using our model; (3) quantifying human disturbance levels, as our pipeline is able to distinguish images containing humans, vehicles and/or domestic animals such as cattle (sheep, cow) and dogs. The current model performances should therefore enable to build reliable metrics estimating human activities along wide gradients.

Importantly, the general high quality of the model results was conserved when applied to an out-of-sample datasets. It is a common error to take model results on validation datasets at face value and assume similar accuracies will be observed in new applications. Previous studies (Whytock et al. 2021) have shown that this is not the case and observed dramatic declines in accuracy when tested on out-of-sample datasets, to the point where the usefulness of the model could sometimes be called into question. Our out-of-sample tests, based on heterogeneous images from new and unseen locations suggests that our model performances are robust to new applications. Additionally, these out-of-sample tests were designed to test the applicability of our model to real-world scenarios. It revealed that using the average prediction score over temporally-close pictures (i.e. sequence) can improve classification results if classification at the sequence-level is sufficient. Our results also show that the common approach of field practitioners to setup camera-traps to take several images at each trigger can be leveraged when using classification models. Our work also highlights that the reported performance of a classification model trained without sequence information is likely an under-estimation of the performance it could achieve at the sequence-level. Whether there are better ways to aggregate image-level scores into sequence-level scores remains to be studied. We also note that sequence or temporal information could be directly integrated into the training step, as done in context CNN models (Beery et al. 2020; Tuia et al. 2022), but we could not use this approach here as many partners who provided images for training the models provided batches of images that did not come from the same sequences.

### 4.2 Lessons learned from a successful multi-partner initiative

One of the main strengths of the DeepFaune initiative lies in the creation of a nation-wide network among key actors in French biodiversity research, conservation and management. Under the lead of an academic research group originating from two distinct laboratories, over 50 partners have shared camera-trap pictures and videos allowing to build what is likely to be one of the largest databases of camera-trap medias in France, in both number (more than 2 millions annotated pictures and 20 thousands of annotated videos), location heterogeneity (e.g. mountains, wetlands).

This success should not hide the technical challenges of working with such a large number of actors. Building the original database clearly revealed the intricacies of dealing with multiple, high-volume data transfers as well as with the strong heterogeneity in data acquisition and organization among the partners. The devil is in the details, and harmonizing directory, file and species names were all very time-consuming tasks that had to be dealt with by combining automatic and manual interventions. Partner-specific data management appears as one of the strongest barriers to creating efficient large-scale datasets that do not depend on a single monitoring program (as opposed to the Serengeti Snapshot (Swanson et al. (2015)) dataset for instance). Interestingly however, some of our individual partners were members of the same institution but worked in different sites. For example, we received data from several national parks that all belong to the same institution (Parc Nationaux de France). Similarly, several teams from the Office Français de la Biodiversité shared data, all having different formats. In such cases, it would seem beneficial for the master institution to provide detailed data management guidelines that would allow standardized data management internally. In this context, centralized data management platforms that enforce data standardization, developed either within institutions or at national or international levels, might facilitate future works.

Having numerous and diverse partners is key however to ground our work in the reality of end-users, as the DeepFaune initiative was originally conceived to develop a free and easy-to-access tool of sufficient quality for field practitioners. Beyond the data collection stage, a regular communication between the leading academic team and the partners was critical to identify key expectations and potential difficulties in appropriation. Expectations shared by all partners were that the machine learning approach should be easy to implement and run on a standard personal computer. This is a long-standing issue regarding deep-learning models, as even when training is not required and only predictions are expected, the installation of libraries necessary for computation (e.g. Tensorflow, PyTorch) can be difficult. Uploading pictures on an online platform running its own servers naturally solves this issue. We however found that many partners were reluctant to follow this approach either because of the need to upload gigabytes of images online or because the data-sharing policies were too stringent. We thus decided to implement our approach in a freely available, cross-platform (Windows, macOS, Linux), Python-based software whose developments can be followed at https://www.deepfaune.cnrs.fr. In general, we feel that the diversity of users and of user needs will prevent finding a one-size-fits-all solution and allow for numerous initiatives to co-exist.

Expectations of our partners differed regarding the classification model and how to use its prediction to sort out images. What makes an ‘optimal’ sorting strategy might in fact vary between partners. Some partners are interested in minimizing false negatives over false positives. This can be the case when studying sensitive species, such as wolves, as positive classifications will in any case be verified manually. For other studies, the balance between false positives and false negatives matters less. This is generally the case in occupancy modeling studies of common species (Gimenez et al. 2022). We also foresee that prediction scores will, in a near future, be directly used in the inference process allowing the uncertainty of the classification to be propagated into the estimation of any metric of interest. Irrespective of whether this will be successful or not, in response to this diversity of needs, we decided to let the users decide of a threshold on the prediction score below which the images will not be classified and requires manual inspection. We believe that this approach gives an important level of flexibility and that users will be able to learn what threshold works best for them by trial-and-error.

## 4.3 Conclusion

In conclusion, our work provides a rather successful classification model of the European fauna in camera-trap images, which can be easily used by practitioners through the Deepfaune software on self-hosted images using a standard personal computer. Such an approach contrasts with some recent developments that favor cloud-computing. Our work however remains a work-in-progress and feedback on the use of the model and the associated software, as well as annotations of images for which the model failed, should allow to improve the model’s performances in a near future.

## Statements and Declarations

### Contributions and disclosure of competing interests

N. Rigoudy and S. Chamaillé-Jammes initiated the project and the collaborative network. V. Miele developed the models and the graphical-user-interface with G. Dussert. G. Dussert performed all the experiments to validate the models’ performance. B. Spataro supervised the use of the CC LBBE/PRABI computing servers for storage and computation. All five make up the DeepFaune team and all contributed to organize the data collection, clean-up and management. E. Chetouane and J. Rabault contributed significantly to the model and software development. Other authors contributed data on their behalf or on behalf of their institutions. The authors declare that they have no financial nor intellectual conflicts of interest with the content of this article.

## Acknowledgments

This work was performed using the computing facilities of the CC LBBE/PRABI. This work has benefited from discussions initiated within meetings of the GDR ‘Ecologie Statistique’ of the CNRS. V. Miele would like to thank the LECA laboratory for having hosted him in Chambéry, and L. Humblot for insightful discussions about the software. Also, for many partners, field technicians and students contributed largely to data collection, and they are acknowledged for their critical work. Some partners could not share images at the time of our study but actively participated in the discussions around future model developments: PNR des Ardennes, PNR de Chartreuse, PNR du Haut-Languedoc, PNR du Haut Jura, PNR du massif des Bauges, PNR de Préalpes d’Azur, PN de la Vanoise and PN des Calanques, FNC.

## Supplementary Information

**Table S1.** Performance of the DeepFaune detector model. The DeepFaune detector model, based on YOLOv8s with a threshold of 0.6 was compared to the MegaDetectorV5a (MDV5a) model with a threshold of 0.5. The thresholds were chosen to optimize the trade-off between accuracy in detecting animals and accuracy in classifying empty images. Animals were classified as small when they are broadly smaller than a fox (e.g. bird, squirrel, lagomorph, mustelide, micromammal). The two models gave similar results, MDV5a being slightly better for images containing larger animals. MDV5 was however much slower (2 to 3 seconds per image) than the DeepFaune detector (0.3 seconds per image).

| Image type | % of images with an animal detected, using MDV5a | % of images with an animal detected, using the DeepFaune model | Average % Intersection-over-Union between MDV5a and the DeepFaune predictions | Number of out-of-sample images |
| --- | --- | --- | --- | --- |
| Large animals | 85.1 | 79.6 | 0.9 | 119831 |
| Small animals (without video frames) | 63.8 (0.77) | 66.2 (0.73) | 0.9 (0.84) | 7036 (1697) |
| Empty images | 2.98 | 2.97 | NA | 8583 |

**Figure S.1.**
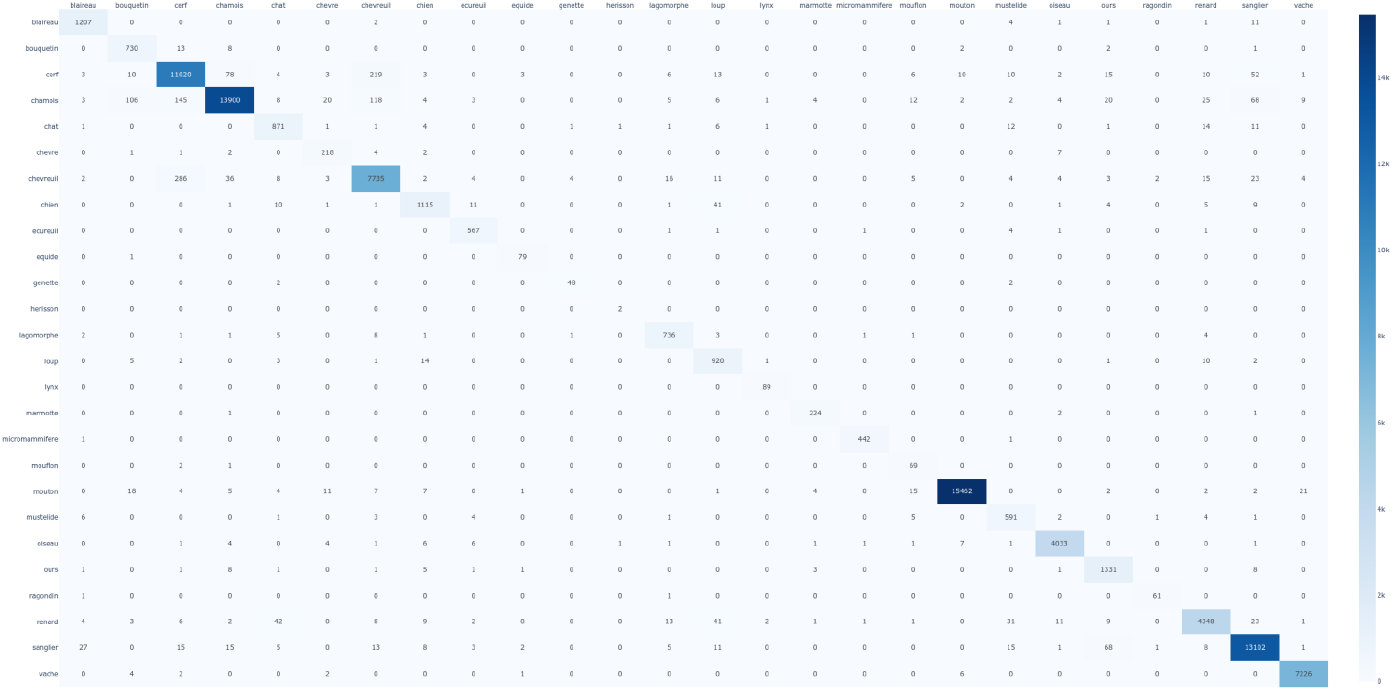
Confusion matrix of the predictions of the CNN-based classifier on the validation dataset. Ground truth is in row, prediction in column.

**Figure S.2.**
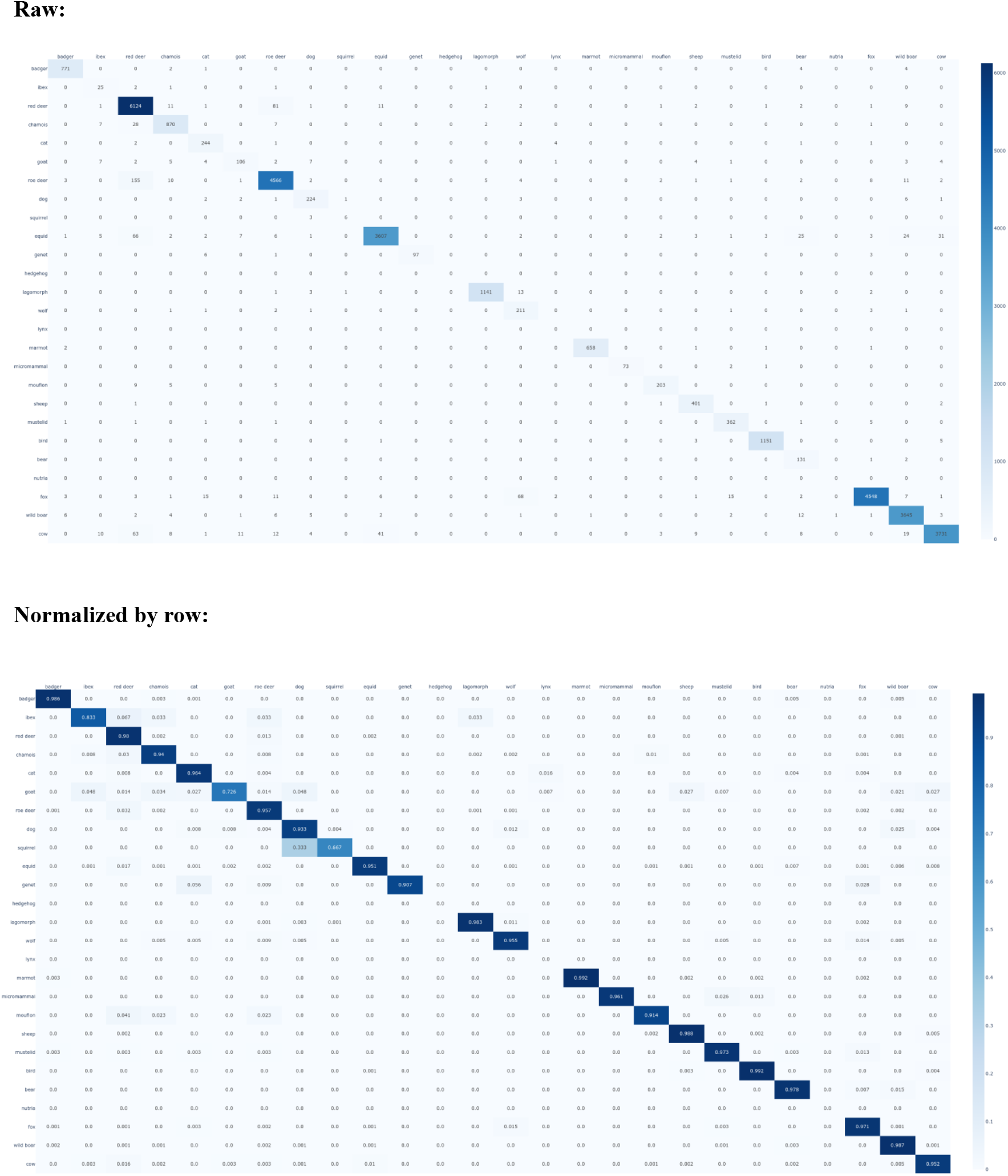
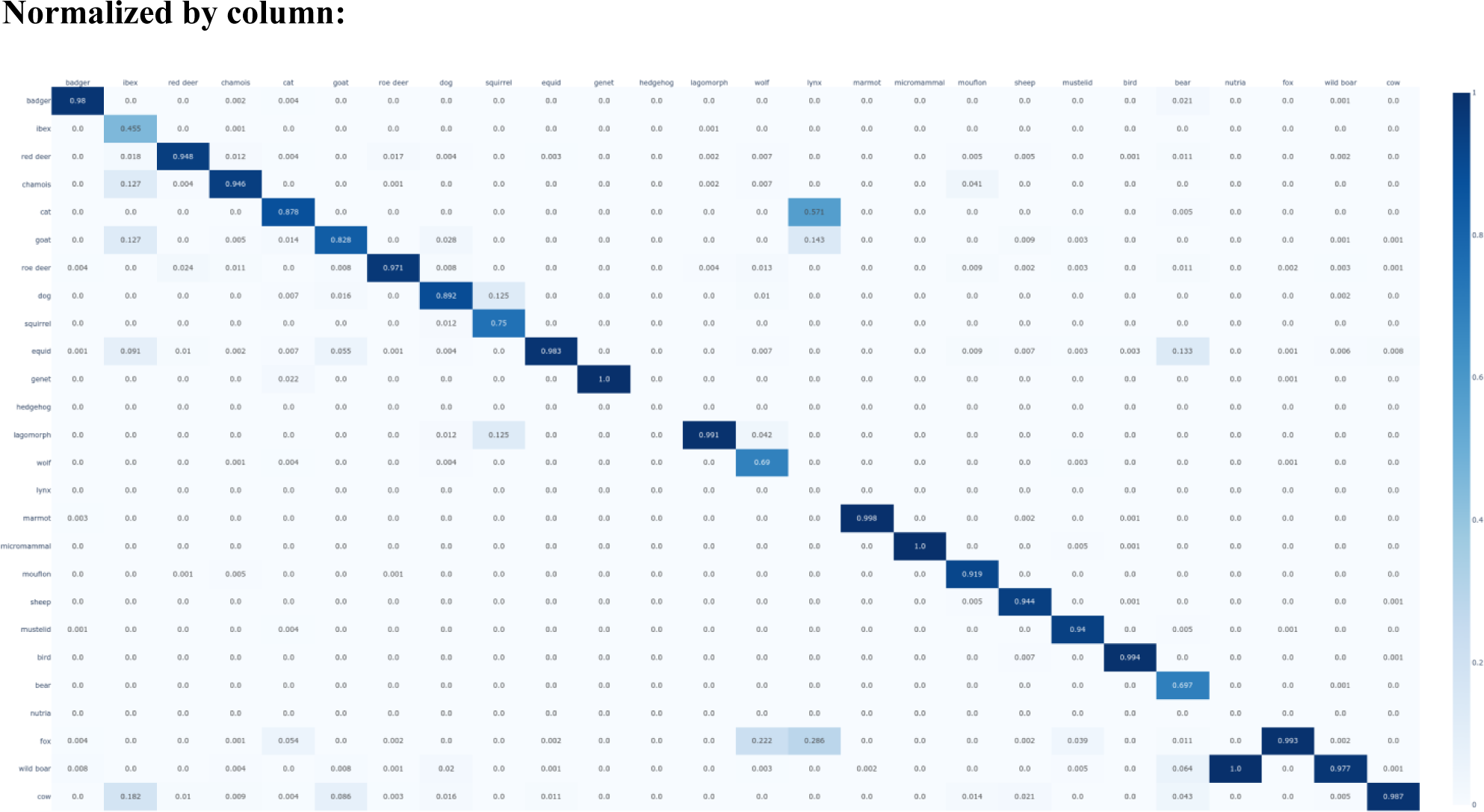
Confusion matrices (raw, normalized by row, or normalized by column, respectively) of the predictions of the CNN-based classifier on the out-of-sample dataset. Ground truth in row, prediction in column.

